# A Point Mutation in the RNA Recognition Motif of *CSTF2* Associated with Intellectual Disability in Humans Causes Defects in 3′ End Processing

**DOI:** 10.1101/2020.01.02.893107

**Authors:** Petar N. Grozdanov, Elahe Masoumzadeh, Vera M. Kalscheuer, Thierry Bienvenu, Pierre Billuart, Marie-Ange Delrue, Michael P. Latham, Clinton C. MacDonald

## Abstract

*CSTF2* encodes an RNA-binding protein that is essential for mRNA cleavage and polyadenylation (C/P). No disease-associated mutations have been described for this gene. Here, we report a mutation in the RNA recognition motif (RRM) of *CSTF2* that changes an aspartic acid at position 50 to alanine (p.D50A), resulting in intellectual disability in male patients. In mice, this mutation was sufficient to alter polyadenylation sites in over 1,000 genes critical for brain development. Using a reporter gene assay, we demonstrated that C/P efficiency of CSTF2^D50A^ was lower than wild type. To account for this, we determined that p.D50A changed locations of amino acid side chains altering RNA binding sites in the RRM. The changes modified the electrostatic potential of the RRM leading to a greater affinity for RNA. These results highlight the importance of 3′ end mRNA processing in correct expression of genes important for brain plasticity and neuronal development.

## INTRODUCTION

Learning, memory, and intelligence require synaptic plasticity involving persistent changes in neural gene expression. Abnormalities in brain development can result in neurodevelopmental disorders that impact intellectual functioning, behavioral deficits involving adaptive behaviors, and autism spectrum disorders. Most neuronal developmental changes are accomplished by post-transcriptional processes that alter the regulation and protein coding capacity of specific mRNAs (Miura et al., 2014; Raj and Blencowe, 2015). Similarly, neurodegenerative conditions are associated with altered mRNA metabolism. Frequently, mRNAs in the brain have extremely long 3′ untranslated regions (UTRs), and altered 3′ UTRs in *MECP2* (Rett syndrome), *APP* (Alzheimer disease), and *HTT* (Huntington disease) are associated with improper metabolism of each of these mRNAs leading to disease states. Though rarer, monogenic forms of intellectual deficiencies offer specific insight into neuronal plasticity and development. The most common monogenic forms of intellectual deficiency are X-linked (XLID, Hu et al., 2016; Jamra, 2018). For example, Fragile X Syndrome is caused by expansion of CGG repeats in the 5′ UTR of *FMR1*, resulting in loss-of-function of FMRP, an RNA-binding protein that promotes transport of mRNAs to dendrites for protein synthesis at synaptic sites (Davis and Broadie, 2017; Mila et al., 2018). Similarly, mutations in mRNA processing genes encoding decapping enzymes (Ahmed et al., 2015; Ng et al., 2015), the polyglutamine binding protein 1 (PQBP1, Kalscheuer et al., 2003), spliceosomal proteins (Carroll et al., 2017), hnRNA-binding proteins (Bain et al., 2016), mRNA surveillance proteins (Tarpey et al., 2007), and cleavage and polyadenylation factors (Gennarino et al., 2015) cause intellectual disabilities. These disorders are associated with changes in the mRNA processing landscape, especially 3′ end cleavage and polyadenylation (C/P), highlighting the importance of RNA processing in controlling neuronal function (Fontes et al., 2017; MacDonald, 2019; Szkop et al., 2017; Wanke et al., 2018).

More than eighty different proteins are involved in mRNA C/P (Shi et al., 2009). Two core C/P factors are the cleavage and polyadenylation specificity factor (CPSF) and the cleavage stimulation factor (CstF). CPSF has six subunits that recognize the polyadenylation signal (AAUAAA and closely related sequences), cleaves the nascent pre-mRNA, then recruits the poly(A) polymerase, which adds up to 250 non-template adenosines to the upstream product of the cleavage reaction (Shi and Manley, 2015; Tian and Manley, 2017). CstF is the regulatory factor in C/P, consisting of three subunits, CstF-50 (gene symbol *CSTF1*), CstF-64 *(CSTF2)*, and CstF-77 *(CSTF3)*. *CSTF2,* the RNA-binding component of CstF, binds to U- or GU-rich sequences downstream of the cleavage site through its RNA recognition motif (RRM, Grozdanov et al., 2018b). As such, *CSTF2* regulates gene expression in immune cells (Chuvpilo et al., 1999; Edwalds-Gilbert and Milcarek, 1995; Takagaki et al., 1996), spermatogenesis (Dass et al., 2007; Hockert et al., 2011; Li et al., 2012), and embryonic stem cell development (Youngblood et al., 2014; Youngblood and MacDonald, 2014). Furthermore, the *CSTF2* paralog, *Cstf2t* has been shown to affect learning and memory in mice (Harris et al., 2016).

Here, we describe members of a family in whom a single nucleotide mutation in the RRM of *CSTF2* changes an aspartic acid at amino acid 50 to an alanine (D50A). The probands presented with non-syndromic intellectual disability. The mutation, which is X-linked, co-segregated with the clinical phenotype. In a reporter gene assay, *CSTF2* containing the D50A mutation *(CSTF2^D50A^)* reduced C/P by 15%. However, the CSTF2^D50A^ RRM showed an almost 2-fold increase in its affinity for RNA. Solution state nuclear magnetic resonance (NMR) studies showed that the overall backbone structure of the CSTF2^D50A^ RRM was similar to that of wild type. However, repositioning of side chains in RNA binding sites and the α4-helix resulted in changes in the electrostatic potential and fast timescale protein dynamics of the RNA-bound state. Together, these changes led to an increased affinity of the CSTF2^D50A^ RRM for RNA due to a faster k_on_ rate. Differential gene expression analysis of RNA isolated from brains of male mice harboring the D50A mutation (*Cstf2^D50A^*) identified fourteen genes important for synapse formation and neuronal development. Genome-wide 3′ end sequencing of polyadenylated mRNAs identified more than 1300 genes with altered C/P sites, leading to alternative polyadenylation of genes involved in neurogenesis, neuronal differentiation and development, and neuronal projection development. Our results indicate that in affected patients, different affinity and rate of binding to the nascent mRNAs of the mutant CSTF2 RRM could misregulate expression of genes critical for brain plasticity and development.

## RESULTS

### Males in Family P167 presented with mild intellectual disability and a mutation in the RNA Recognition Motif in *CSTF2*

Family P167 is of Algerian origin currently living in France. The family has four affected male individuals (II:1, II:2, III:1, and III:2, Figure 1) in two different generations connected through the maternal germline. This pattern of inheritance suggested an X-linked trait carried by females I:2 and II:4. The index proband, a 6-year-old boy (III.1) was presented to the Division of Pediatrics at Bicêtre Hospital (Kremlin-Bicêtre, France) with developmental delays. There was no parental consanguinity. He has one affected brother (II.2) and two affected maternal uncles (I.3 and I.4). All affected males were born after uneventful pregnancies and delivery. The index proband (III.1) was born at term. Weight (3,760 g), height (50 cm), and occipitofrontal circumference (OFC, 34 cm) were normal at birth. Motor milestones were reached within normal limits. He walked at 14.5 months. Developmental delay was recognized in early childhood during the first years at school and speech development was retarded. Differential scales of intellectual efficiency (Échelles Différentielles d’Efficience Intellectuelle, EDEI, Fiasse and Nader-Grosbois, 2012) showed low verbal and nonverbal communication skills, with no other consistent clinical phenotype. He was not institutionalized and attended a school for children with additional speech therapy and special assistance. Brain MRI showed no specific abnormalities except for an abnormal hypersignal at the posterior thalamic nuclei due to an episode of intracranial hypertension at the age of 9.5 years (data not shown). His two uncles (ages 44- and 46-years-old, II.1 and II.2, Figure 1A) attended a special school for children with learning difficulties. They are both married and employed; one uncle has a 15-year-old unaffected girl, and the other a 7-year-old unaffected girl and a 5-year-old unaffected boy.

**Figure 1.**
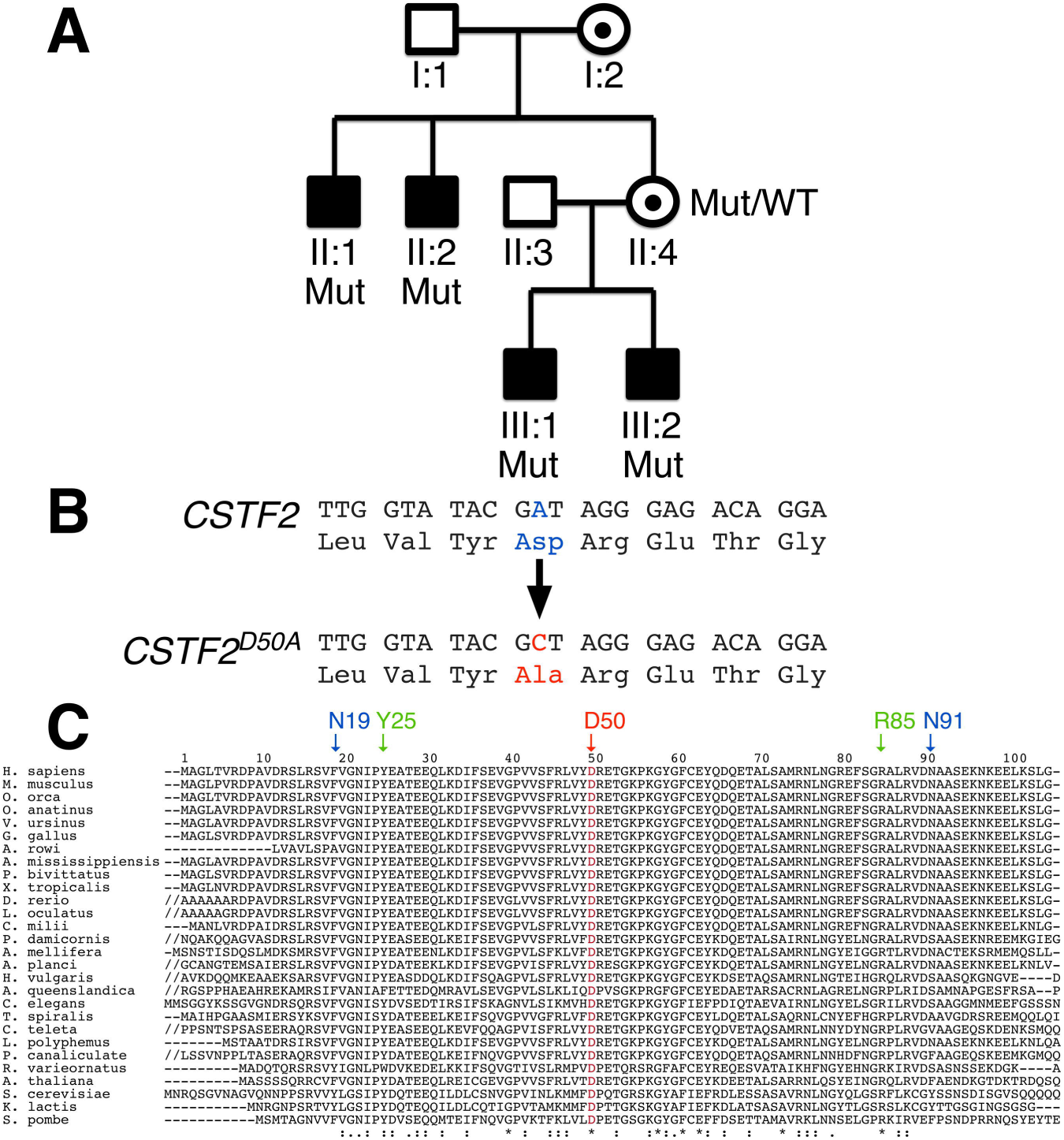
Family P167 carries a mutation in the X-linked *CSTF2* gene that affects intelligence in males. (A) Pedigree of family 167 showing the index proband (III:1) who was 6 years old at the time of the study, his affected brother (III:2), and two affected uncles (II:1 and II:2). Females (I:2 and II:4) were carriers for the trait, consistent with an X-linked recessive trait. Individuals labeled Mut (II:1, II:2, III:1, III:2) or Mut/WT (II:4) were tested for co-segregation of the mutation with the clinical phenotype by PCR and Sanger sequencing of the specific products. (B) The missense mutation in exon 3 of *CSTF2* substitutes an alanine codon for an aspartic acid at position 50 (p.D50A) within the RNA recognition motif (RRM). (C) CLUSTAL alignment of *CSTF2* orthologs in twenty-eight species indicate that the aspartic acid (D) at position 50 (red arrow) is highly conserved. Amino acids comprising site-I (Y25 and R85) and site-II (N19 and N91) are indicated in green and blue, respectively.

Initial karyotype analysis testing for Fragile X and a mutation search in a few selected XLID genes, including *SLC6A8*, *NLGN4* and *JARID1C* gave normal results. X chromosome exome sequencing of the index proband (III:1) as part of a large study of more than 500 unrelated males from families with likely XLID (Hu et al., 2016) identified a 3 bp deletion in *STARD8*, and single nucleotide variants in *ALG13*, *COL4A5* and *CSTF2*. All variants co-segregated with the phenotype: they were present in all affected males and were transmitted through the maternal germline (Figure 1A and data not shown). The single amino acid deletion identified in STARD8 is present in 180 control males in the Genome Aggregation Database (gnomAD, non-neuro control individuals, https://gnomad.broadinstitute.org/) (Lek et al., 2016), and was therefore interpreted as a benign polymorphism. *ALG13* is an established epilepsy-associated gene in females (Bissar-Tadmouri et al., 2014), but the *ALG13* missense mutation in Family P167 (chrX:110951509G>A, NCBI RefSeq NM_001099922.3:c.638G>A, p.S213N, rs374748006) has been reported in four control males (gnomAD, non-neuro) and was predicted as a polymorphism or benign variant by MutationTaster2 (Schwarz et al., 2014) and the ClinVar database (Landrum et al., 2016). Mutations in *COL4A5* have been associated with X-linked Alport syndrome, also known as hereditary nephritis. The *COL4A5* single nucleotide mutation (chrX:107846262C>A, NM_000495.4:c.2215C>A, NP_203699.1:p.P739H) has not been reported in any publicly available database. However, the same proline residue substituted by an alanine (p.P739A) is most likely a benign polymorphism (44 hemizygous males in gnomAD).

In contrast, the *CSTF2* missense mutation (exon 3, chrX:100077251A>C (GRCh38.p12), RefSeq NM_001306206.1:c.149A>C) was identified as novel and was not reported in more than 14,000 control individuals from publicly available databases including gnomAD. The mutation was predicted as disease causing or pathogenic by several prediction tools, including MutationTaster2 (disease causing), DANN (pathogenicity score of 0.9956 with a value of 1 given to the variants predicted to the most damaging, Quang et al., 2015) and a high CADD score of 26.9 (Rentzsch et al., 2019). Furthermore, at the gene level, *CSTF2* is highly constrained with zero known loss-of-function mutations and a Z-score of 3.43 (ratio of observed to expected = 0.37) for missense mutations in gnomAD. Conceptual translation of the cDNA demonstrated a mutation of aspartic acid (GAT) to alanine (GCT) within the RNA recognition motif (RRM) of *CSTF2* (p.D50A, Figure 1B). The aspartic acid at position 50 is conserved in every *CSTF2* ortholog surveyed (Figure 1C). We designated this allele *CSTF2^D50A^*.

### Polyadenylation Efficiency is Reduced by the *CSTF2^D50A^* Mutation

Previously, we showed that mutations in the RRM of *CSTF2* affected RNA binding and altered C/P efficiency using the Stem-Loop Assay for Polyadenylation (SLAP, Grozdanov et al., 2018b; Hockert et al., 2010). To determine the effectiveness of *CSTF2^D50A^* in control of C/P, we performed SLAP using CSTF2 constructs with an N-terminal MS2 coat protein domain (MCP-CSTF2 and MCP-CSTF2^D50A^). Wild type MCP-CSTF2 produced a SLAP value of 4.49 ± 0.16 normalized luciferase units (NLU), whereas MCP-CSTF2^D50A^ consistently achieved 15% lower polyadenylation efficiency (3.92 ± 0.13 NLU) than wild type MCP-CSTF2 (Figure 2A).

**Figure 2.**
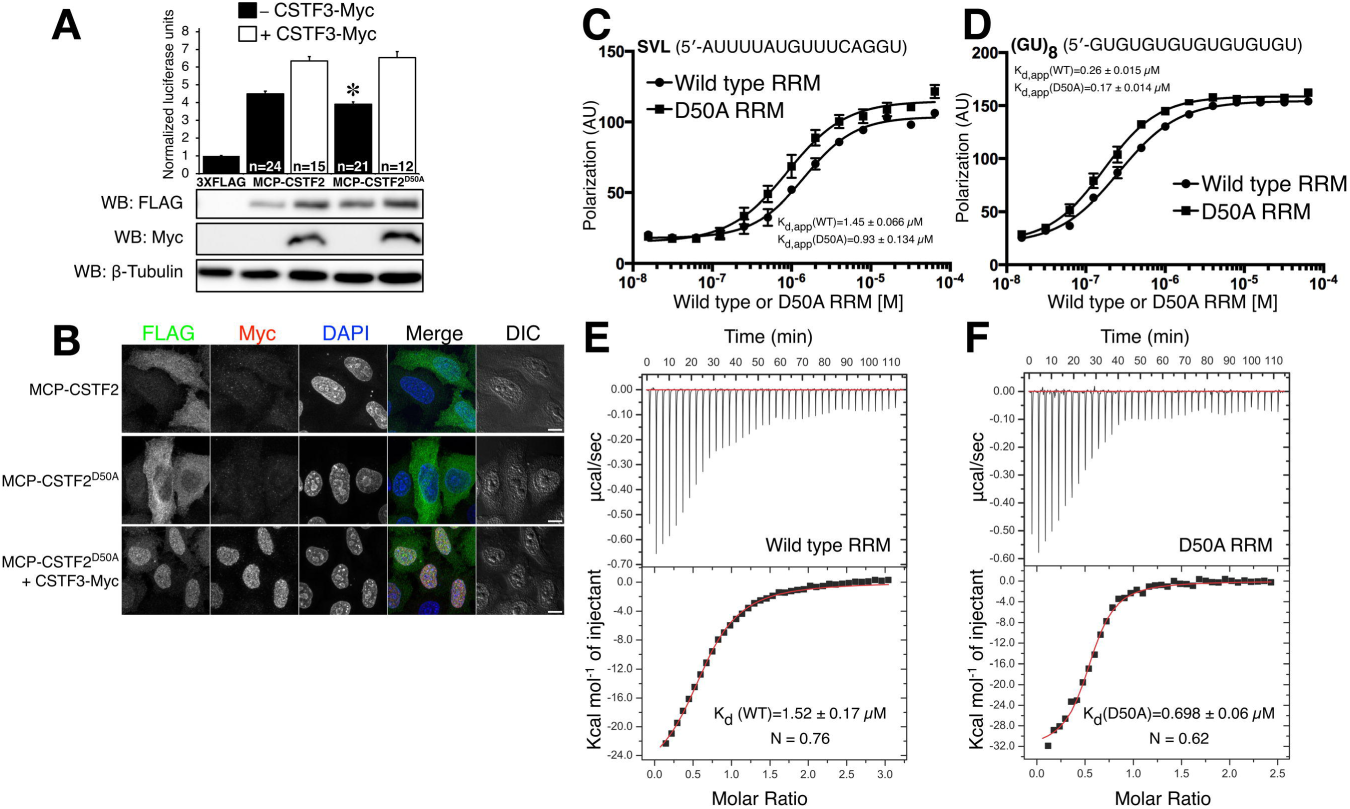
*CSTF2^D50A^* is less efficient for C/P because it binds substrate RNA with a higher affinity. (A) SLAP results showing normalized luciferase units (NLU) in HeLa cells without MCP-CSTF2 (–), in cells transfected with MCP-CSTF2 or MCP-CSTF2^D50A^ (black bars) and with CSTF3-Myc (white bars). Western blots to show the expression of FLAG-tagged MCP-CstF-64 (WB: FLAG) or Myc-tagged CSTF3-Myc (WB: Myc). [β-tubulin was used as a loading control. (B) Immunofluorescent images of the described constructs stained with antibodies against the FLAG tag for MCP-CSTF2 (green), Myc tag for CSTF3 (red), and counterstained with DAPI to delineate the nucleus. (C) The K_d_ for RNA binding was determined for the isolated RRM domain of wild type CSTF2 and CSTF2^D50A^ mutant via the change in fluorescence polarization of a 3′-end labeled RNA substrate (C, D) and isothermal titration calorimetry (ITC; E, F). Changes in fluorescence polarization for wild type and D50A RRMs binding to SVL (C) and (GU)_8_ (D) substrate RNAs. ITC thermograms of wild type (E) and D50A (F) RRM binding to SVL RNA. The K_d_ and the number of binding sites (N) are indicated on the figures along with the corresponding standard deviation from three replicates. Raw injection heats are shown in the upper panels and the corresponding integrated heat changes are shown in the bottom panels versus the molar ratio of RNA to protein. K_d_s and thermodynamics, derived from the fits of the ITC data, are provided in Table S1.

Mutations in the RNA binding site I and II of CSTF2 affect the synergetic function of CSTF2 and CSTF3 (Grozdanov et al., 2018b). Therefore, we co-expressed CSTF3 with either MCP-CSTF2 or MCP-CSTF2^D50A^ to determine whether the D50A mutation affects cleavage and polyadenylation in the presence of CSTF3. CSTF3 increased SLAP for both wild type and MCP-CSTF2^D50A^ to a similar extent, effectively eliminating the reduction observed in the MCP-CSTF2^D50A^ construct expressed alone. Mutations in the *CSTF2* RRM altered nuclear-to-cytoplasmic localization (Grozdanov et al., 2018b). Therefore, we hypothesized that the *CSTF2^D50A^* mutation might also affect the ratio between nuclear and cytoplasmic *CSTF2.* Immunohistochemical staining revealed that CSTF2^D50A^ was localized more in the cytoplasm than wild type CSTF2 (Figure 2B). Co-expression of *CSTF3* with *CSTF2^D50A^* increased the nuclear localization of CSTF2^D50A^ similar to wild type CSTF2 protein (Figure 2B and Grozdanov et al., 2018b).

### The CSTF2^D50A^ RRM Has a Greater Affinity for RNA

Because the aspartic acid (D) to alanine (A) mutation changes the charge in the loop connecting the [β2 and [β3-strands (Figure 4 and Figure S1, Deka et al., 2005; Grozdanov et al., 2018b), we wanted to determine whether RNA binding was altered in the *CSTF2^D50A^* RRM mutant. To measure binding via fluorescence polarization/anisotropy, bacterially expressed CSTF2 and CSTF2^D50A^ RRMs (amino acids 1–107, Grozdanov et al., 2018b) were incubated with fluorescently-labeled RNA oligonucleotides from either the SV40 late transcription unit (SVL, MacDonald et al., 1994) or (GU)_8_ (Deka et al., 2005). The wild type RRM bound to the SVL and (GU)_8_ RNAs with K_d, app_ of 1.45 ± 0.07 µM and 0.26 ± 0.02 µM, respectively (Figure 2C, D and Table S1). The CSTF2^D50A^ mutant RRM bound the two RNAs with significantly higher affinities (K_d, app_ 0.93 ± 0.13 µM and 0.17 ± 0.01 µM for the SVL and (GU)_8_ RNA, respectively, Figure 2C, D, Table S1).

To confirm the RNA-binding affinities, we performed isothermal titration calorimetry (ITC) titrations using the SVL RNA oligonucleotide. The average K_d_ was 1.52 ± 0.17 µM for wild type RRM and 0.70 ± 0.06 µM for the CSTF2^D50A^ RRM (Figure 2E and F, Table S1), which were in the same range as the K_d_s measured before. In both cases, binding was driven by a favorable enthalpy overcoming an unfavorable entropy. Yet, the enthalpy and entropy for mutant and wild type binding to SVL RNA were different: the ΔH for wild type binding to RNA was −24.6 ± 1.64 kcal mol^-1^ with a ΔS = −55.5 ± 5.44 cal mol^-1^ K^-1^ (Figure 2E, Table S1); whereas, the D50A mutant had a ΔH = −32.4 ± 0.33 kcal mol^-1^ and ΔS = −79.7 ± 0.89 cal mol^-1^ K^-1^ for binding (Figure 2F, Table S1). Thus, the enthalpy-entropy compensation for the D50A mutant binding to SVL RNA was greater than that of the wild type RRM, leading to the higher observed affinity, suggesting that CSTF2^D50A^ would bind to RNAs with a greater affinity during C/P.

### The D50A Mutation Affects the Side Chain Interactions and Electrostatic Surface of the RRM

To characterize the dynamic changes that occur in the CSTF2^D50A^ RRM upon binding to RNA, we turned to solution state nuclear magnetic resonance (NMR) spectroscopy, characterizing the fingerprint of the general backbone conformation via 2D ^15^N,^1^H heteronuclear single quantum coherence (HSQC) spectra for the wild type and mutant RRMs. Predictably, the overlay of the HSQC spectra showed differences in the peak positions of the residues in the loop where the D50A mutation is located (Figure 3A and C, green bars). Otherwise, the majority of the residues in the CSTF2^D50A^ RRM showed almost identical NMR spectra as the wild type.

**Figure 3.**
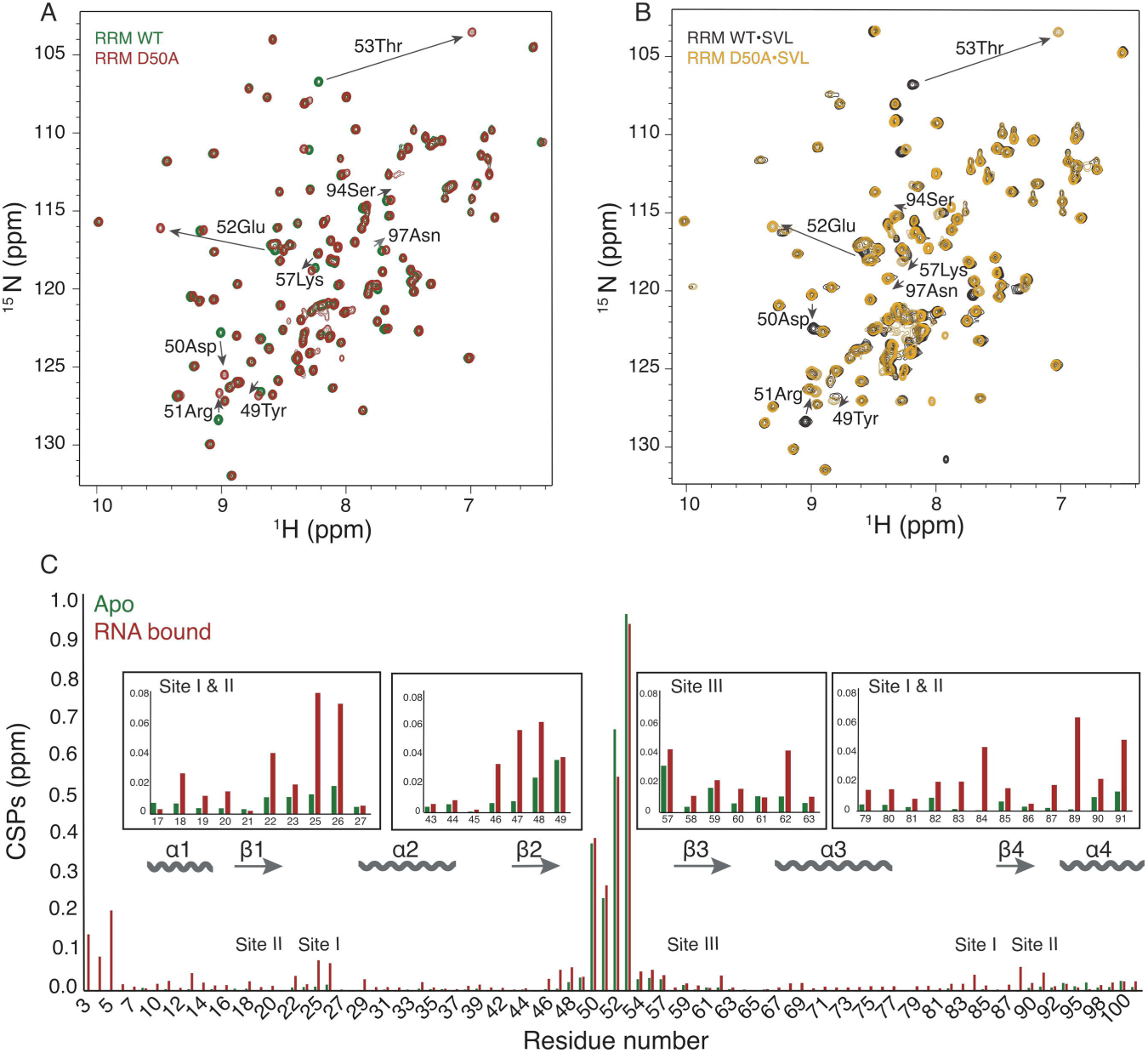
The p.D50A mutation perturbs the environment of the β-sheet and C-terminal α-helix. The 2D ^15^N, ^1^H HSQC of apo (A) RRM WT (green) and RRM D50A (red) and RNA-bound (B) RRM WT•SVL (black) and RRM D50A•SVL (yellow) complexes. Data were collected at 600 MHz and 27 °C. (C) Bar graph indicating backbone amide chemical shift perturbations (CSP) for the RNA-free (Apo) WT and D50A RRM (green) and SVL RNA-bound WT and D50A mutant RRM (red). The CSPs are calculated from the HSQC spectra from panels A and B.

**Figure 4.**
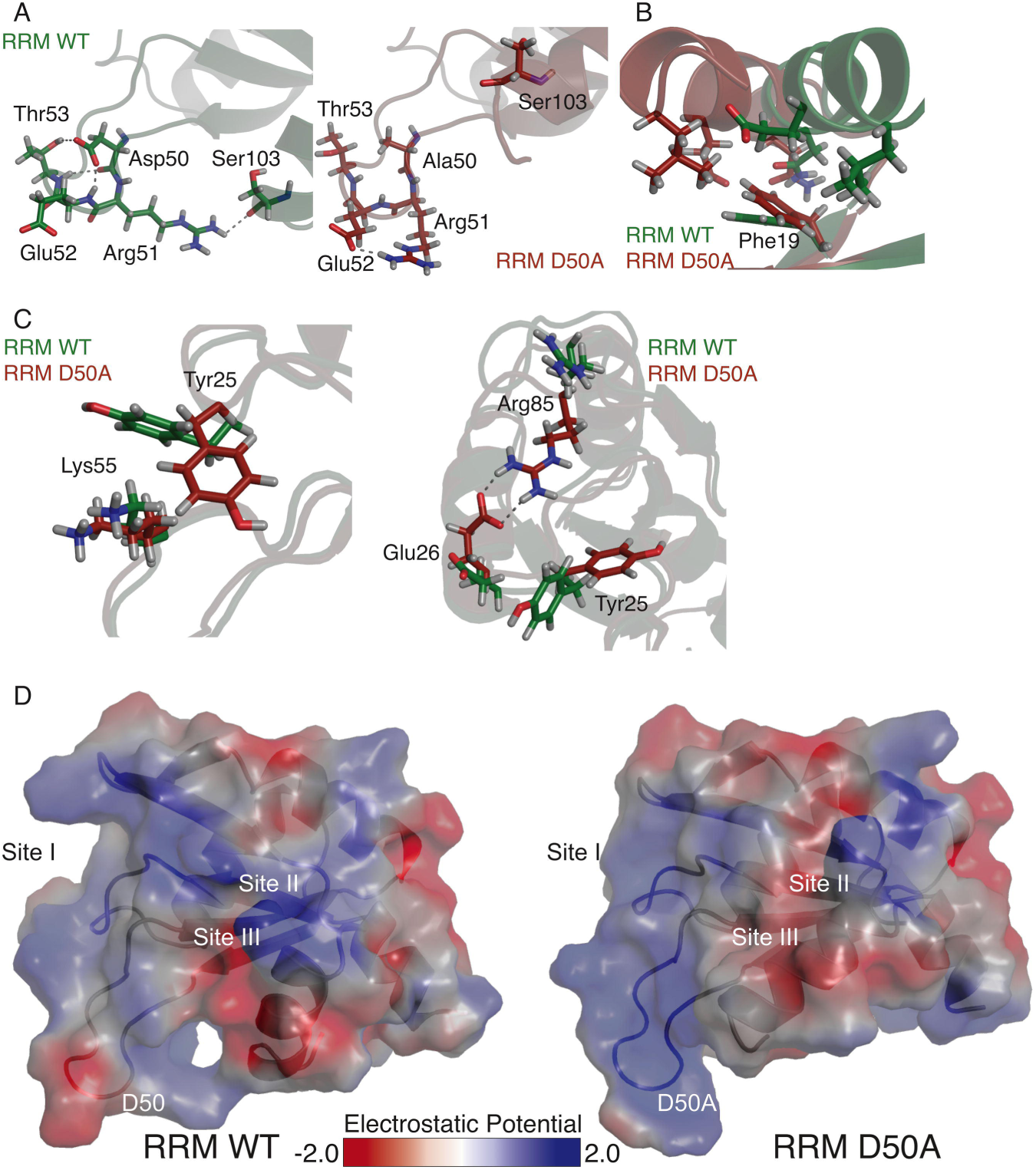
The altered side chain interactions of CSTF2^D50A^ lead to different local electrostatic surface potentials. Panels A–C show the different side chain orientations and interactions present in the wild type CSTF2 RRM (green) and mutant CSTF^D50A^ RRM (red). See Figure S1 for the alignment of the overall structures. (D) Calculated electrostatic potential for wild type (left) and D50A mutant (right) RRM. The red-to-blue surface representation highlights negative-to-positive electrostatic potential from −2 to 2 k_b_T/e, where k_b_ is Boltzmann’s constant, T is the temperature in Kelvin, and e is the charge of an electron.

Next, we determined the three-dimensional structures of the CSTF2 and CSTF2^D50A^ RRMs using CS-Rosetta (Ramelot et al., 2009; Shen et al., 2008). The ^1^HN, ^15^N, ^13^Cα, and ^13^C[3 backbone chemical shifts were submitted along with amide ^1^HN-^15^N residual dipolar coupling (RDC) data collected in the presence of Pf1 bacteriophage (Hansen et al., 1998). Resulting structures were rescored with PALES against an independent set of amide RDC data collected in the presence of DMPC/DHPC bicelles (Wang et al., 1998; Zweckstetter, 2008). In the absence of RNA, the CSTF2^D50A^ RRM was almost identical to the CSTF2 RRM structure (Figure S1A; 3.081 Å all atom root-mean-square deviation, RMSD), consistent with circular dichroism data (Figure S2A). The major difference between the RRMs was observed in the α4-helix (Figure S1A), which angled away to make room for the repositioned sidechains of the β4-strand to interact with the α4-helix. A small difference was also noted in the relative twist of the β-sheet (Figure S1B), which includes RNA binding sites-II and -III.

These differences in secondary structural elements resulted from repositioning of the side chains for the residues in the loop surrounding the D50A mutation and in the amino acids involved in site-I (Tyr25 and Arg85) and -II (Phe19 and Asn91; Figure 1C and 4A–C), which are two of the three sites identified as important for RNA interactions in *Rna15,* the yeast homolog of *CSTF2* (Pancevac et al., 2010). Specifically, in wild type CSTF2, the carboxylic acid side chain of Asp50 formed a hydrogen bond with the hydroxyl side chain of Thr53 (Figure 4A, left). Replacing the hydrogen bond acceptor with a methyl group disrupted this interaction in CSTF2^D50A^ (Figure 4A, right). In addition, the Arg51 side chain formed a new ion pair interaction with the side chain carbonyl group of Glu52, instead of the backbone carbonyl of the terminal helix α4 residue Ser103, leading to re-orientation of helix α4 (Figure 4A). In the CSTF2^D50A^ RRM, the β1-strand had a different twist, which allows the aromatic ring of Phe19 (site-II, Figure 1C) to lift up towards the RNA binding pocket (Figure 4B), which also contributed to the reorientation of helix α4 in the mutant as the edge of the Phe19 aromatic ring now packs against the aliphatic portions of Asn97, Glu100, and Leu104 of helix α4. In binding site-I of the wild type RRM, the Tyr25 aromatic ring was adjacent to the amino side chain on Lys55, forming a π-cation interaction (Figure 4C). However, in CSTF2^D50A^, the Tyr25 ring moved away, allowing Glu26 to form a hydrogen bond with Arg85 (site-I residue), increasing the rigidity in binding site-I (Figure 4C right). Thus, local differences in side chain arrangement, particularly in binding sites-I and -II, contribute to the differences we observed in RNA binding.

To determine whether the electrostatic surface potentials of the RRMs also contributed, we calculated the potentials using the Adaptive Poisson-Boltzmann Solver (Baker et al., 2001). Changing the negatively charged Asp50 to the non-charged Ala altered the charge distribution of the loop from negative to positive in the mutant (Figure 4D). Binding site-I, which forms upon RNA binding (Pancevac et al., 2010) in the wild type, was within a cavity with low positive electrostatic potential for RNA binding (Figure 4D, left). However, in the D50A mutant, the hydrogen bonding between Arg85 and Glu26 (Figure 4D) moved the positively charged side chain of Arg85 towards the inside of cavity, resulting in a higher positive electrostatic potential distribution in the site-I (Figure 4D, right). We hypothesize that this larger positive electrostatic potential could be the driving force for the more favorable enthalpy of binding and the tighter RNA binding affinity.

### The D50A Mutation Affects the Structure and Dynamics of the RNA-bound RRM

To test effects of RNA binding, the CSTF2 and CSTF2^D50A^ RRMs were titrated with SVL RNA to saturation (molar ratio of 2.3:1, RNA to protein) with changes monitored in 2D ^15^N, ^1^H HSQC spectra (Figures 3B and 5A). Both proteins showed the same binding patterns (i.e., direction, magnitude, and exchange regime) for most residues (e.g., Val18, Val20, Ala27, Asp90, and Leu47; Figure 5A). Gly54 located within the D50 loop was perturbed upon addition of RNA to the CSTF2^D50A^ RRM but not the wild type RRM (Figure 5A), indicating a potentially new interaction in the mutated domain. Next, we calculated amide chemical shift perturbations (CSPs) between SVL-bound CSTF2 and CSTF^D50A^ RRMs (Figure 3C, red bars). The largest differences were observed in the loop containing the mutation, similar to unbound RRMs (Figure 3C, green bars), and other significant amide CSPs (i.e., above the mean, 0.032 ppm for apo and 0.048 ppm for RNA-bound) were observed in the RNA-bound domain for the N-terminus (residue 5 to 7) near binding site-I (25 and 85), site-II (residues 19 and 91; Figure 3C), and in the β2-strand preceding the mutation (Figure 3B and C, red bars).

**Figure 5.**
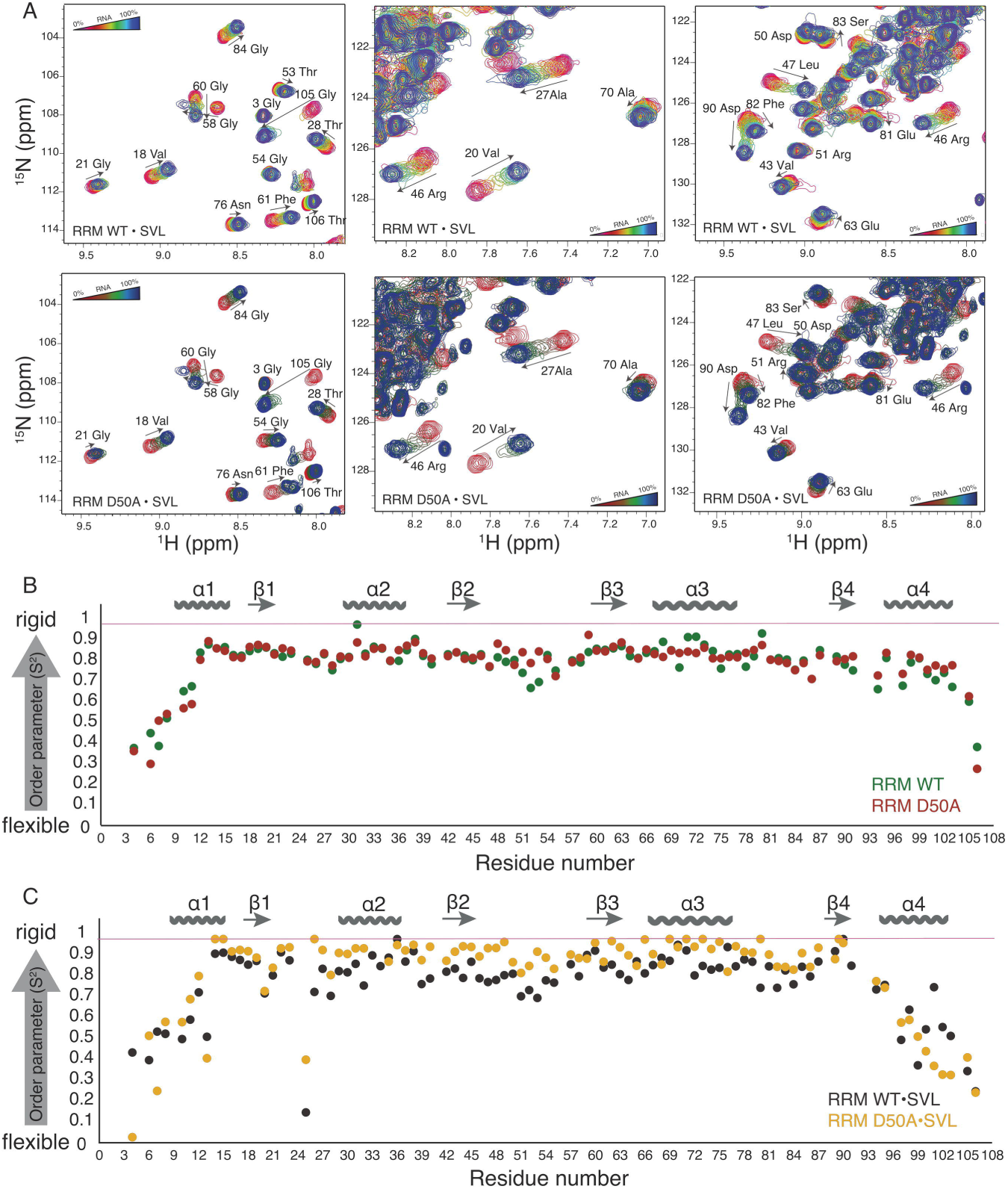
The p.D50A mutation affects the RNA-binding kinetics and dynamics of the RRM. (A) Overlays of the 2D ^15^N, ^1^H HSQC spectra for the titration of CSTF2 (top panels) and CSTF2^D50A^ (bottom panels) RRMs. Red-yellow-blue and red-green-blue gradients represent the 0% to 100% titration of wild type and D50A RRMs with SVL RNA, respectively. Titration data were acquired at 600 MHz and 27 °C. All spectra are shown with the same contour base level relative to noise. The magnitude and direction of each titrated residue is shown with arrows and assignment. (B and C) Backbone amide ^15^N order parameter (S^2^) for (B) apo RRM WT (green) and RRM D50A (red) and (C) SVL RNA-bound RRM WT•SVL (black), and D50A•SVL (yellow). The pink dashed lines at an S^2^ value of 1.0 denotes the maximum value for the order parameter.

The on- and off-rates of RNA binding were determined with a simple two state ligand binding model using the 2D NMR line shape analysis program (TITAN; Figure S3, Waudby et al., 2016). From the RNA-induced perturbations that were in the fast and intermediate exchange regimes, we calculated a K_d_ of 0.70 ± 0.02 µM for wild type and 0.54 ± 0.02 µM for CSTF2^D50A^, which followed the trend established by fluorescence polarization and ITC (Table S1). The off-rates (k_off_) were similar for wild type (305.5 ± 1.4 s^-1^) and CSTF2^D50A^ (312.4 ± 2 s^-1^) RRMs. Both on-rates were the same as or exceeded the rate of diffusion, a feature seen in other protein-nucleic acid complexes where electrostatic attraction accelerates the on-rate (Riggs et al., 1970; von Hippel and Berg, 1989). However, the on-rate (k_on_) for CSTF2^D50A^ (5.83 ± 0.21 x 10^8^ M^-1^ s^-1^) was faster than the rate for wild type (4.35 ± 0.12 x 10^8^ M^-1^ s^-1^). We conclude, therefore, that the lower K_d_ for CSTF2^D50A^ was primarily due to the faster on-rate for RNA binding to D50A compared to the wild type.

We examined ^15^N R_1_, R_2_, and nuclear Overhauser effect (NOE) relaxation values at 600 MHz, 27 °C using the model-free approach to derive backbone order parameters (S^2^), which report on the amplitude of fast timescale motion and the global correlation time (τ_c_) (Lipari and Szabo, 1982a, b). In the absence of RNA, the S^2^ values for the CSTF2^D50A^ RRM domain differed from wild type only in the D50A loop, which became more rigid. The average S^2^ for the entire domain was 0.83 for both (Figure 5B and Figure S4A; root mean square deviation, RMSD = 0.049). However, for the RNA-bound structures, the S^2^ values indicated different amide group flexibilities in the mutant RRM (average S^2^ were 0.78 and 0.84 for wild type and mutant RRM, respectively, Figure 5C and Figure S4B; RMSD = 0.12). SVL RNA binding to the CSTF2 RRM reduced S^2^ values throughout the β-sheet (e.g., residues 18–22 and 44–48), indicating increased flexibility upon RNA binding, in agreement with earlier studies (Deka et al., 2005; Perez Canadillas and Varani, 2003). However, SVL RNA binding to the CSTF2^D50A^ RRM increased S^2^ values for the β-sheet and the α3-helix adjacent to site-II (Phe19 and Asn91, Figure 5C). This indicated greater rigidity in the pico-to-nanosecond time scale in the RNA binding surface of the D50A mutant when bound to RNA. The order parameter is a measure of conformational entropy within a protein (Wand, 2013; Yang and Kay, 1996). Indeed, the wild type RRM, which became more flexible upon RNA binding, pays less of an entropic penalty compared to the D50A mutant (Table S1), which maintained the same flexibility upon RNA binding. Thus, the overall tertiary fold of the CSTF2^D50A^ RRM was nearly identical to wild type while unbound to RNA.

We also determined whether the point mutation in D50A changed the stability of the secondary structure of the motif. Using CD spectroscopy, we determined the changes in ellipticity at 216 nm (α-helix) and 222 nm (β-sheet) upon thermal and chemical denaturation (Figure S2B and C). Both RRMs were thermally stable with minimal changes in ellipticity at 100 °C (data not shown). Therefore, we used increasing concentrations of guanidine to probe the differences in RRM stability. The inflection point, where half of the protein was denatured, for the wild type protein was reached at 2 M guanidine and was decreased to at 1.5 M for the CSTF2^D50A^ RRM mutant. This result suggested that secondary structure of the mutant was less stable than the wild type.

### The D50A Mutation Causes Differential Gene Expression in Mice

To examine effects of the D50A mutation *in vivo*, we developed *Cstf2^D50A^* mice using CRISPR-Cas9 technology. Hemizygous male (*Cstf2^D50A/Y^*) and homozygous female (*Cstf2^D50A/D50A^*) mice are viable and fertile. Male *Cstf2^D50A/Y^* mice appear to be runted and may have other musculoskeletal abnormalities that will be described elsewhere (K. A. White, P. N. Grozdanov, and C. C. MacDonald, in preparation). We isolated whole brain RNA from 50-day old male wild type and *Cstf2^D50A/Y^* littermates and performed RNA-seq. Only fourteen genes showed significant differential mRNA expression between wild type and *Cstf2^D50A/Y^* male mice; eleven were up-regulated and three were down-regulated (Figure 6A). Functional gene enrichment analysis for biological processes (Liao et al., 2019) indicated that up-regulated differentially expressed genes were involved in several brain developmental pathways including hindbrain development, neural tube patterning, head and brain development (Figure 6B), demonstrating that the D50A mutation in *Cstf2* influenced overall expression of key genes in mice.

**Figure 6.**
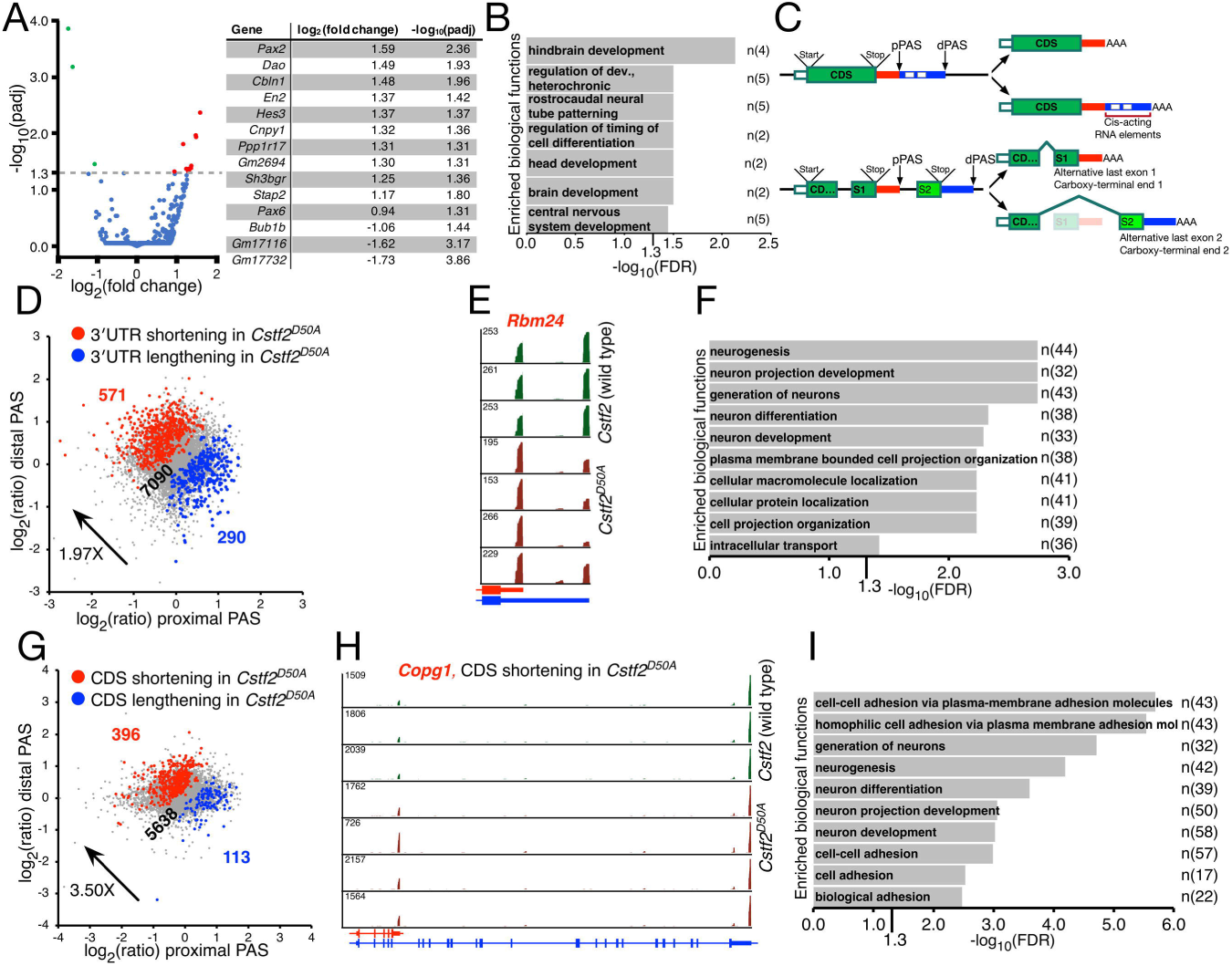
Gene expression and 3′ APA analyses in brains of *Cstf2^D50A/Y^* mice. (A) Differential gene expression analysis of RNA-seq from total brain samples isolated from 50-day old male wild type and *Cstf2^D50A/Y^* mice (three wild type and five mutants). Left, volcano plot of the differentially expressed genes. Right, list of genes that are differentially expressed, fold change (log_2_) and adjusted p value (-log_10_) are indicated. (B) Gene ontology for biological functions of differentially expressed genes in the brains of wild type and *Cstf2^D50A/Y^* mice. Number (n) on the right indicates the number of genes found in each functional category. (C) Schematic illustration of APA events analyzed. (Top) Genes in which APA changes the length of the last (3′-most) exon. (Bottom) Genes in which APA occurs in the coding region (CDS) changes the protein coding potential of the gene. Proximal polyadenylation site (PAS), distal PAS, start and stop codons in translation, and regions containing cis-acting RNA elements (e.g., miR and RNA-binding protein binding sites) are indicated. (D) Scatter plot showing expression change of proximal PAS isoform (x-axis) and that of the distal PAS isoform (y-axis) in total RNA. Genes with significantly shortened or lengthened 3′ UTRs (p < 0.05, Fisher’s exact test) in *Cstf2^D50A/Y^* total mouse brains (three wild type and five mutant) are highlighted in red and blue, respectively. Color coded numbers indicated the number of genes shortened, lengthened, or showing no change. (E) C/P in the last exon of the *Rbm24* gene in wild type (green) or *Cstf2^D50A/Y^* (rust) mouse brains switch from the distal to the proximal poly(A) sites. Proximal and distal PASs are indicated with the maximum RPM values shown. (F) Top ten gene ontology categories (FDR<0.05) of enriched biological functions in *Cstf2^D50A/Y^* mice showing lengthening of their 3′-most exons. Number of genes (n) that are detected in each functional category are shown on the right. (G) Scatter plot showing expression change of proximal PAS isoform (x-axis) and that of distal PAS isoform (y-axis) in total RNA. Genes with significantly shortened or lengthened CDSs (p < 0.05, Fisher’s exact test) in *Cstf2^D50A/Y^* or wild type mouse brains are highlighted in red and blue, respectively. Color coded numbers indicated the number of genes shortened, lengthened, or showing no change. (H) C/P in the *Copg1* gene shortens the CDS in *Cstf2^D50A/Y^* mouse brains (rust) compared to wild type (green). Maximum RPM values are shown for each animal. Gene structure and different transcripts are shown at the bottom. (I) Top ten gene ontology categories (FDR<0.05) of enriched biological functions in *Cstf2^D50A/Y^* brains showing shortened transcripts in their CDS. Number of genes (n) that are detected in each functional category are shown on the right.

### The D50A mutation Affects Alternative Polyadenylation (APA) of over One Thousand Genes in Mice

To examine the changes in C/P in the *Cstf2^D50A/Y^* mice, we performed genome-wide 3′ end sequencing (Li et al., 2016; Li et al., 2015) on total RNA isolated from brains from 50-day old male wild type and *Cstf2^D50A/Y^* littermates. To simplify the analysis, we focused on *(i)* genes in which APA changed the length of the last (3′-most) exon, and *(ii)* genes in which APA caused changes in the coding region (CDS, Figure 6C). The site that was farther from the transcription start site of the gene was designated the distal polyadenylation site (dPAS) and the nearer one was designated the proximal polyadenylation site (pPAS) for both types of APA changes (Figure 6C).

In total, 1370 genes changed polyadenylation sites in *Cstf2^D50A/Y^* mouse brains compared to wild type. The number of the genes showing shortening of the last exon in the D50A mice was 1.97-fold greater than those showing the opposite pattern (571 vs. 290, Figure 6D and Table S2). An example of last exon shortening is the RNA Binding Motif Protein 24 *(Rbm24)* gene (Figure 6E). Gene ontology (GO) analysis of the genes shortening the last exon showed localization of macromolecules in cells, retrograde transport, cell migration, and more (Table S3). GO terms of the genes lengthening the last exon indicated enrichment in neurogenesis, neuron development, and more (Figure 6F).

The number of genes in which APA caused changes in the CDS in *Cstf2^D50A/Y^* animals was 3.50-fold greater than the number of genes showing the opposite pattern (396 vs. 113, Figure 6G and Table S4). For example, the mRNA for the shorter version of the Coatomer Protein Complex Subunit Gamma 1 *(Copg1)* gene was increased in *Cstf2^D50A/Y^* mouse brains (Figure 6H). This isoform was previously reported to be enriched in granule cells of the brain (Jereb et al., 2018). GO term analysis for the 113 genes lengthening their CDSs did not reveal ontologies that were enriched (FDR<0.05, TableS2). However, GO terms for genes with shorter CDSs were enriched for neurogenesis, neuro development and cell adhesion (Figure 6I). We propose that in the patients carrying the D50A mutation, the balance of the critical protein isoforms involved in brain development is disrupted, contributing to their cognitive disability.

## DISCUSSION

Many monogenic intellectual deficiencies are X-linked (XLIDs) because of hemizygosity of X-chromosomal genes in males (Hu et al., 2018; Penrose, 1938; Raymond, 2006). The X-linked *CSTF2* gene is essential for embryonic growth and development (Youngblood et al., 2014; Youngblood and MacDonald, 2014). Therefore, it was surprising that a single nucleotide mutation in *CSTF2* would result in structural and functional changes in 3′ end mRNA processing yet not have lethal effects. Other mutations in the RRM of *CSTF2* have been reported, but with no associated disease. For example, gnomAD (Karczewski et al., 2019) reports a substitution of a threonine at position 53 with an isoleucine (three heterozygous females and one hemizygous male out of 182,201 alleles) and substitution of a tyrosine at position 59 with a cysteine (one heterozygous female out of 183,168 alleles); both were aphenotypic. As reported here (Figure 1), it appears that the *CSTF2^D50A^* mutation affects primarily intellectual functions and speech development in the affected males, but effects on other physiological functions were not noted, suggesting that the brain is more sensitive to this particular mutation than other organs. We propose, therefore, that the p.D50A mutation in CSTF2 results in non-syndromic intellectual deficiency in males by altering RNA binding during C/P, thus changing the expression of key genes in neurodevelopment.

Deleterious mutations in essential genes are rare because loss of their functions generally result in embryonic lethality (Gluecksohn-Waelsch, 1963). Only one mutation involving a core polyadenylation protein, *NUDT21* (which encodes the 25 kDa subunit of cleavage factor I_m_), has been implicated in neuropsychiatric disorders (Gennarino et al., 2015). In this case, duplications of *NUDT21* result in altered polyadenylation of *MECP2,* reducing its translation and resulting in Rett syndrome-like symptoms. We did not observe altered expression of *Mecp2* in our mouse model (not shown), but observed altered sites of polyadenylation in many other neurodevelopmental genes (Figure 6).

RNA-contact residues in the CSTF2 RRM play specific roles during C/P (Grozdanov et al., 2018b). Introduction of the D50A mutation into CSTF2 resulted in a small but consistent reduction in C/P efficiency (Figure 2A). Such a reduction in activity would likely result in altered C/P in vivo, favoring proximal sites over more distal sites (Hwang et al., 2016; Yao et al., 2012). Analysis of genome-wide polyadenylation changes in the *Cstf2^D50A/Y^* mice indicated exactly that: shorter RNAs (3′ most exons and changes in the last exon) were enriched in brains of *Cstf2^D50A/Y^* mice (Figure 6D, G). Residues within the CSTF2 RRM are also important for functional interactions between CSTF2 and CSTF3 during C/P (Grozdanov et al., 2018b). However, we did not observe a reduction in the ratio between MCP-CSTF2^D50A^ alone and MCP-CSTF2^D50A^ co-expressed with CSTF3, which we previously observed with CSTF2 RRM binding site-I and -II mutants (Figure 2). This suggests that the D50A mutation causes the phenotype independent of the interactions with CTSF3. Furthermore, the small decrease in the polyadenylation efficiency in our reporter system might suggest that the mutated protein is retained longer in the cytoplasm (Figure 2B), possibly through increased interaction with cytoplasmic RNAs (Grozdanov et al., 2018b).

With reduced C/P efficiency, we observed an increased affinity of the CSTF2^D50A^ RRM for RNA (Figures 3 and 5), probably also contributing to the retention of CSTF2^D50A^ in the cytoplasm (Figure 2B). The K_d_ of the CSTF2^D50A^ RRM for RNA was less than half that of wild type CSTF2 due to the faster k_on_ rate (Table S1). Based on the on- and off-rates, the D50A mutant binds to RNA faster than wild type but releases the RNA with the same off-rate. It has been previously shown that the β2-β3 loop (D50A loop here) is important for the shape recognition of RNA in some RRM containing proteins (Skrisovska et al., 2007; Stefl et al., 2005). Our structures for the CSTF2 and CSTF2^D50A^ RRMs allowed us to model the differences in RNA binding based on the relative orientation of the side chains of the mutant RRM, electrostatic potential, and rigidity of individual backbone atoms of the β-sheet, causing faster RNA binding in the mutant (Figure 4D) by helping to overcome the greater entropic penalty of binding resulting from the enhanced rigidity of the β-sheet in the mutant.

To confirm that the CSTF2^D50A^ mutation had a neurophysiological effect, we created *Cstf2^D50A^* mice with the same mutation. Several labs have examined effects of knockdowns of *CSTF2* in human cells in culture, but noted altered C/P in only a relatively small number of genes (Gruber et al., 2012; Kim et al., 2010; Martin et al., 2012; Yao et al., 2012). Unlike those studies, we observed C/P site changes in 1370 genes in the brains of hemizygous *Cstf2^D50A/Y^* mice (Figure 6). The majority of the changes in our study favored proximal PASs, effectively shortening the CDSs or shortening the length of the 3′ UTRs. Why did the knockdown studies have fewer consequences than the D50A point mutation? Possibly, their results were confounded by the presence of τCstF-64 (gene symbol *CSTF2T)* compensating for decreased CSTF2 in those cells. In support of that idea, knockout of *Cstf2t* in mice resulted in few phenotypes in most cells (where *Cstf2* was also expressed), but led to severe disruption of spermatogenesis in germ cells (where *Cstf2t* was expressed in the absence of *Cstf2)* (Dass et al., 2007).

But it is also possible that cells in the brain (either neurons, glia, or both) are more sensitive to mutations in *CSTF2.* It has long been known that the brain uses alternative polyadenylation extensively to provide transcriptomic diversity and mRNA targeting signals for plasticity and behavioral adaptation (Fasken and Corbett, 2016; Miura et al., 2013; Raj and Blencowe, 2015; Zhang et al., 2005). By extending occupancy times on the pre-mRNA, the D50A mutation likely changes the balance of APA to favor shorter mRNAs, thus reducing the effectiveness of over one thousand mRNA transcripts (Figure 6). The brain also expresses a neuron-specific alternatively-spliced isoform of *CSTF2* that contains up to 49 extra amino acids (Shankarling et al., 2009; Shankarling and MacDonald, 2013). Thus, we speculate that the D50A mutation might interact with the additional domain in [βCstF-64 in an as-yet undefined manner to give the brain phenotype.

How does an increase in the affinity of CSTF2 for RNA result in altered cleavage and polyadenylation? Previous work showed that formation of the C/P complex takes 10–20 seconds for a weak site; assembly on stronger sites is faster (Chao et al., 1999). We speculate that the increased affinity of CSTF2^D50A^ alters the rate of formation of the CstF complex on the downstream sequence element of the nascent pre-mRNA during transcription (MacDonald et al., 1994). We further speculate that the specificity of binding is reduced in CSTF2^D50A^. This combination of faster binding and reduced specificity could promote increased C/P of weaker sites, which tend to be more proximal (Martin et al., 2012), not unlike the effects of slower RNA polymerase II elongation (Liu et al., 2017; Pinto et al., 2011). Comparable issues may arise with CSTF2’s role in histone mRNA processing efficiency, affecting the cell cycle (Romeo et al., 2014; Youngblood et al., 2014). These C/P changes subsequently affect post-transcriptional regulation of key mRNAs by changing 3′ UTR regulation by revealing miRNA or RNA-binding protein sites, or by changing targeted localization of mRNAs to neural projections (Ciolli Mattioli et al., 2019; Fontes et al., 2017; Hafner et al., 2019; Jereb et al., 2018; Nazim et al., 2017; Taliaferro et al., 2016; Wanke et al., 2018).

Finally, we note that *CSTF2T,* the testis-expressed paralog of *CSTF2* (Dass et al., 2002), is primarily associated with male infertility (Dass et al., 2007; Hockert et al., 2011; Tardif et al., 2010). However, female but not male *Cstf2t^-/-^* mice also showed reduced spatial learning and memory (Harris et al., 2016). *Cstf2t* has been implicated in the control of global gene expression (Li et al., 2012), splicing (Grozdanov et al., 2016), small nuclear RNA expression (Kargapolova et al., 2017), and histone gene expression (Grozdanov et al., 2018a; Youngblood et al., 2014) in male germ cells. Mutations in *CSTF2* may have even more striking effects in target tissues. Studying these mutation-induced changes in gene expression will be important for understanding the mechanisms of polyadenylation as well as the intricacies of neuronal development.

## EXPERIMENTAL PROCEDURES

### Human Subjects

All cases had a normal karyotype, were negative for FMR1 repeat expansion, and large insertions or deletions were excluded using array Comparative Genomic Hybridization (CGH). The study was approved by all institutional review boards of the participating institutions collecting the samples, and written informed consent was obtained from all participants or their legal guardians.

### DNA isolation and X-chromosome Exome Sequencing and Segregation Analysis

DNAs from the family members were isolated from peripheral blood using standard techniques. X-chromosome exome enrichment using DNA from the index patient, sequencing and analysis was performed as previously described (Hu et al., 2016). Segregation analysis of variants of uncertain clinical significance was performed by PCR using gene-specific primers flanking the respective variant identified followed by Sanger sequencing.

### Animal Use and Generation of *Cstf2^D50A^* Mice

All animal treatments and tissues obtained in the study were performed according to protocols approved by the Institutional Animal Care and Use Committee at the Texas Tech University Health Sciences Center in accordance with the National Institutes of Health animal welfare guidelines. TTUHSC’s vivarium is AAALAC-certified and has a 12/12-hour light/dark cycle with temperature and relative humidity of 20–22 °C and 30–50%, respectively.

B6;*Cstf2^em1Ccma-D50A^* founder mice (herein *Cstf2^D50A^)* were generated by Cyagen US Inc. (Santa Clara, CA). To create C57BL/6 mice with a point mutation (D50A) in the *Cstf2* locus, exon 3 in the mouse *Cstf2* gene (GenBank accession number: NM_133196.6; Ensembl: ENSMUSG00000031256) located on mouse chromosome X was selected as the target site. The D50A (GAT to GCT, see Figure 1) mutation site in the donor oligo was introduced into exon 3 by homology-directed repair. A gRNA targeting vector and donor oligo (with targeting sequence, flanked by 120 bp homologous sequences combined on both sides) was designed. *Cas9* mRNA, sgRNA and donor oligo were co-injected into zygotes for KI mouse production. The pups were genotyped by PCR, followed by sequence analysis. Positive founders were bred to the next generation (F1) and subsequently genotyped by PCR and DNA sequencing analysis. Mutants were maintained as a congenic strain by backcrossing four generations to C57BL/6NCrl (Charles River) and subsequently breeding exclusively within the colony. At the time of this study, mice were bred to approximately 10 generations.

### Genotyping of *Cstf2^D50A^* Mice by PCR and Restriction Enzyme Digestion

Genomic DNA was extracted from tail snips of *Cstf2^D50A^* mice by proteinase K digestion followed by isopropanol precipitation (Dass et al., 2007). PCRs were performed using specific primers surrounding the D50A mutation site. The presence of the mutation converted a CGATAGG sequences to CGCTAGG, thus introducing a *Bfa*I restriction site. Digestion of the PCR products with *Bfa*I revealed the presence of the mutation.

### Cell culture, Transfection, Stem-Loop Assay for Polyadenylation, immunohistochemistry and Western Blots

Culturing of HeLa cells was performed as described (Grozdanov et al., 2018b). Transfection was carried out in a 24-well plates using lipofectamine LTX (ThermoFisher Scientific) and 250 ng of a mixture of plasmid DNAs. Luciferase measurements were performed between 36-48 hours after the transfection (Grozdanov et al., 2018b; Hockert and Macdonald, 2014). Western blots to assess the abundance of the proteins were performed on the same volumes of lysates obtained from the luciferase measurements using the passive lysis buffer supplied with the Dual-Luciferase Reporter Assay System (Promega). Antibodies used for western blots were previously described (Grozdanov et al., 2018b; Wallace et al., 1999). Immunohistochemistry and microscopic imaging were performed as described (Grozdanov et al., 2018b).

### Plasmids and site-directed mutagenesis

The pGL3 plasmid (Promega), Renilla-luciferase construct (SL-Luc) containing a modified C/P site by the addition of two MS2 stem-loop downstream sequences was previously described (Hockert et al., 2010; Maciolek and McNally, 2008). The D50A mutant was created through site-directed mutagenesis (New England Biolabs). The RRM (amino acids 1–107) and D50A mutant RRM was cloned in a bacterial expression vector as a fusion with a His-tag followed by a TEV site at the amino terminal end of the RRM and was previously described (Grozdanov et al., 2018b). All plasmids were verified by sequencing before use.

### Bacterial protein expression and purification

Expression and purification of the proteins over metal affinity resin and His-tag removal was as done before (Grozdanov et al., 2018b). For the NMR experiments, transfected Rosetta (DE3) pLysS cells were grown in 2× minimal M9 media using ^15^NH_4_Cl (1 g/L) and unlabeled or uniformly ^13^C-labeled D-glucose (3 g/L) as sole nitrogen and carbon sources, respectively (Azatian et al., 2019). Induction of the transfected cells with 0.5 mM isopropyl-β-d-thiogalactopyranoside (IPTG), harvest, and purification of the labeled proteins was also carried out as previously described (Grozdanov et al., 2018b).

### Circular dichroism and stability assays

Circular dichroism experiments were performed on a JASCO J-815 instrument in 10 mM sodium phosphate, pH 7.25 with 10 µM of either wild type or D50A RRM protein. The spectra were scanned from 185 to 260 nm with 0.1 nm resolution. The average spectrum was obtained from three technical replicates.

Guanidine-HCl experiments were performed in 5 mM HEPES pH 7.4, 50 mM NaCl with 5 µM protein samples. Guanidine-HCl concentrations ranged from 0.5 to 3 M. Protein samples without guanidine-HCl were used as reference. Spectra were collected between 205 and 230 nm with 0.1 nm resolution. Values for 216 nm and 222 nm were plotted to represent the denaturation of the secondary structure of the proteins for [β-sheets and α-helixes, respectively. The average spectra were obtained from three technical replicates.

### 3′-end fluorescent RNA labeling and fluorescence polarization/anisotropy

SVL (5′-AUUUUAUGUUUCAGGU-3′) and (GU)_8_ (5′-GUGUGUGUGUGUGUGU-3′) RNAs were commercially synthesized (SigmaAldrich). 3′-end fluorescent labeling of the RNAs with fluorescein-5-thiosemicarbazide (ThermoFisher Scientific) was done as previously reported (Grozdanov and Stocco, 2012). The labeled RNAs were used as 3.2 nM final concentration in polarization assays.

Fluorescence polarization experiments were performed in binding buffer (16 mM HEPES pH 7.4, 40 mM NaCl, 0.008% (vol/vol) IGEPAL CA630, 5 μg/ml heparin, and 8 μg/ml yeast tRNA). Purified wild type and D50A RRMs were diluted in 20 mM HEPES pH 7.4, 50 mM NaCl, 0.01% NaN_3_ and 0.0001% IGEPAL CA630 to 32 µM concentration and used for two-fold serial dilutions. The highest final concentration of the protein in the polarization assay was 16 µM. Fluorescence polarization samples were equilibrated for 2 h at room temperature in 96-well black plates (Greiner Bio) and measured on Infinite M1000 PRO instrument (Tecan Inc) with excitation set at 470 nm (5 nm bandwidth) and emission 520 nm (5 nm bandwidth). At least three technical replicates were performed. The apparent dissociation constants were calculated by fitting the data to a modified version of the Hill equation (Pagano et al., 2011) using GraphPad Prism version 5.2 software for Windows (GraphPad Software). Unpaired t-test was performed using Microsoft Excel (Microsoft Corp).

### Isothermal titration calorimetry (ITC)

The RRM wild type, D50A mutant RRM proteins with SVL RNA were dialyzed overnight into 10 mM sodium phosphate, 1 mM tris(2-carboxyethyl)phosphine (TCEP), 0.05% v/v sodium azide, pH 6 using a 2 kDa MWCO dialysis unit (ThermoFisher). Heats of binding were measured using a MicroCal iTC200 calorimeter (GE Healthcare) with a stirring rate of 1000 rpm at 27 °C. For all titrations, isotherms were corrected by subtracting the heats of RNA dilution. The concentrations of proteins and RNA were calculated by absorbance spectroscopy with extinction coefficients of ε_280_ = 5960 M^-1^ cm^-1^ and ε_260_ = 165.7 M^-1^ cm^-1^, respectively. Data were analyzed by Origin 7, and all measurements were performed at least three times.

### NMR techniques

NMR experiments were recorded in 10 mM phosphate buffer pH 6.0 with 1 mM TCEP, 0.05% w/v sodium azide, 0.1 mg/mL 4-(2-aminoethyl)benzenesulfonyl fluoride (AEBSF), and 10% D_2_O using an Agilent 600 MHz (14.1 T) DD2 NMR spectrometer equipped with a room temperature HCN z-axis gradient probe. Data were processed with NMRPipe/NMRDraw (Delaglio et al., 1995) and analyzed with CCPN Analysis (Vranken et al., 2005). Amide chemical shift perturbations (CSPs) were calculated as backbone ^13^Cα, ^13^Cβ, ^13^C′, ^15^N, and ^1^HN resonance assignments of RRM wild type and D50A mutant, both in the apo and SVL-bound states obtained from standard gradient-selected, triple-resonance HNCACB, HN(CO)CACB, HNCO, HN(CA)CO (Muhandiram and Kay, 1994) and CHH-TOCSY (Bax et al., 1990) at 27 °C. Assignment data were collected with a random nonuniform sampling scheme and processed by Sparse Multidimensional Iterative Lineshape-Enhanced (SMILE) algorithm (Ying et al., 2017). Amide group residual dipolar couplings (RDCs; ^1^D_NH_) were measured from 2D ^15^N, ^1^H In-Phase/Anti-Phase (IPAP) experiments recorded on wild type and D50A mutant protein samples in the absence (^1^J_NH_) and presence (^1^J_NH_ + ^1^D_NH_) of filamentous Pf1 bacteriophage (Hansen et al., 1998) and bicelles alignment media (Wang et al., 1998). For samples in Pf1 bacteriophage, a concentrated stock of protein was mixed with 50 mg/mL bacteriophage stock solutions (Asla) to a final concentration of 15 mg/mL. The bicelle samples contained 5% w/v total DLPC and CHAPSO with a molar ratio of 4.2:1 in a 50 mM phosphate buffer, pH 6.8, 50 mM KCl, 1 mM TCEP, 0.05% w/v sodium azide, 0.1 mg/mL AEBSF and 10% D2O. Bicelles were prepared by dissolving DLPC and CHAPSO in phosphate buffer and vortexing for 1 min. Lipids were temperature cycled three times (30 min at 4 °C followed by 30 min at 40 °C), then diluted 1:1 with wild type or D50A RRM. Samples were equilibrated at 27 °C for 1 h before recording the NMR spectra. The DLPC/CHAPSO bicelle samples were stable at 27 °C for three days.

Wild type (BMRB id: 30652) and D50A (BMRB id: 30653) RRM backbone assignments and amide RDC data from Pf1 bacteriophage were submitted to the CS-ROSETTA webserver (https://csrosetta.bmrb.wisc.edu/csrosetta/submit) for structure calculation (Shen et al., 2008). The ten lowest energy structures from CS-ROSETTA were fitted to the RDC data collected in DLPC/CHAPSO bicelles with the PALES program (Zweckstetter and Bax, 2000). The best wild type and D50A mutant structures are available at protein data bank (PDB codes: 6Q2I for wild type and 6TZE for D50A). The surface electrostatic potential for each structure was calculated with the Adaptive Poisson-Boltzmann Solver (APBS), and figures were generated with PyMol (DeLano, 2002).

RNA titration experiments were performed by adding unlabeled SVL RNA to the ^15^N-labeled proteins and monitoring the change in amide chemical shifts in 2D ^15^N, ^1^H HSQC spectra until complete saturation. Binding affinities were calculated from spectra using a two-state ligand binding model in the TITAN program (Waudby et al., 2016). RNA binding on- and off-rates were calculated for both wild type and D50A RRM from amide peaks in the fast and intermediate exchange regimes by TITAN. Backbone ^15^N R_1_ (longitudinal spin-lattice relaxation rate), R_2_ (transverse spin-spin relaxation rate) and heteronuclear {^1^H}-^15^N NOE relaxation data (Kay et al., 1992; Kay et al., 1989) for RRM wild type and D50A mutant in the absence and presence of SVL RNA were acquired at 600 MHz and 27 °C. ^15^N R_1_, R_2_. NOE values were used to calculate the backbone order parameters (S^2^) and the global correlation time (τ_c_) from the model-free approach (Lipari and Szabo, 1982a, b) using “model 2” in modelfree v4.2 (http://nysbc.org/departments/nmr/relaxation-and-dynamics-nysbc/software/modelfree/).

### RNA-seq and 3′-seq

Total brain tissues were collected from three wild type and five mutant 50-day old male animals from the same litter. Brains were immediately treated with TRIzol reagent (ThermoFisher Scientific). RNA isolation, RNA-seq library preparation and high-throughput sequencing (pair end 150-nucleotide sequencing, PE150) and DESeq2 analysis (Love et al., 2014) were performed by Novogene Corporation. Over Representation Enrichment Analysis of the differentially expressed genes were assessed using WebGestalt (Liao et al., 2019) with settings: Mus musculus; Overrepresentation Enrichment Analysis; Geneontology; Biological Process. Reference Gene List was set on genome. Cellular Component and Molecular Function did not return any gene ontology terms with a false discovery rate (FDR) smaller than or equal to 0.05.

3′ end sequencing and library preparation were performed by Admera Health Inc. 3′-seq libraries were prepared using the QuantSeq 3’ mRNA-Seq Library Prep Kit REV for Illumina from Lexogen. Libraries were sequenced with a specific sequencing primer as per the manufacturer recommendations. At least 40M reads were obtained per sample. The sequences were aligned using STAR aligner (Dobin et al., 2013). Aligned reads were normalized and the genes changing the C/P sites were identified. Analysis of APA was performed by Admera Health. Over Representation Enrichment Analysis of the genes changing C/P sites were assessed using WebGestalt (Liao et al., 2019) with settings: Mus musculus; Overrepresentation Enrichment Analysis; Geneontology; Biological Process. Reference Gene List was set on genome; top 10 gene ontology terms with a false discovery rate (FDR) smaller than or equal to 0.05.

## ACKNOWLEDGEMENTS

The authors wish to acknowledge R. Bryan Sutton, Michaela Jansen, Kerri A. White, Charles Faust, Anne Rice, and the members of the Latham laboratory for technical and intellectual contributions. We thank Admera Health for 3′-seq services and Ruijia Wang, Bin Tian, and Yun Zhao for assistance with 3′-seq analysis. Microscopy was performed in the Image Analysis Core Facility supported in part by TTUHSC. We thank Helene Maurey (Bicêtre), Florence Pinton (Bicetre), Brigitte Simon-Bouy (CH Versailles), and Cecile Zordan (Bordeaux), who performed genetic counselling for this family. The authors would also like to thank the Genome Aggregation Database (gnomAD) and the groups who provided exome and genome variant data to this resource. A full list of contributing groups can be found at https://gnomad.broadinstitute.org/about. Support was from the EU FP7 project GENCODYS, grant number 241995, the 2017–2018 Presidents’ Collaborative Research Initiative of Texas Tech University System, the Department of Cell Biology & Biochemistry (Texas Tech University Health Sciences Center), the South Plains Foundation, and the National Institute of General Medical Sciences, and NIH grant 1R35GM128906 (MPL).

## AUTHOR CONTRIBUTIONS

Conceptualization, V.M.K., P.N.G. and C.C.M.; Methodology, V.M.K., P.N.G., E.M. and M.P.L.; Formal analysis, V.M.K., P.N.G., E.M., T.B., P. B., M-A.D., and M.P.L.; Writing—Original Draft, P.N.G.; Writing—Review and Editing, V.M.K., P.N.G., E.M., M.P.L. and C.C.M.; Supervision, P.N.G. and C.C.M.

