## Supplementary figures and images for "A Point Mutation in the RNA Recognition Motif of *CSTF2* Associated with Intellectual Disability in Humans Causes Defects in 3′ End Processing"

### Supplemental Figure S1

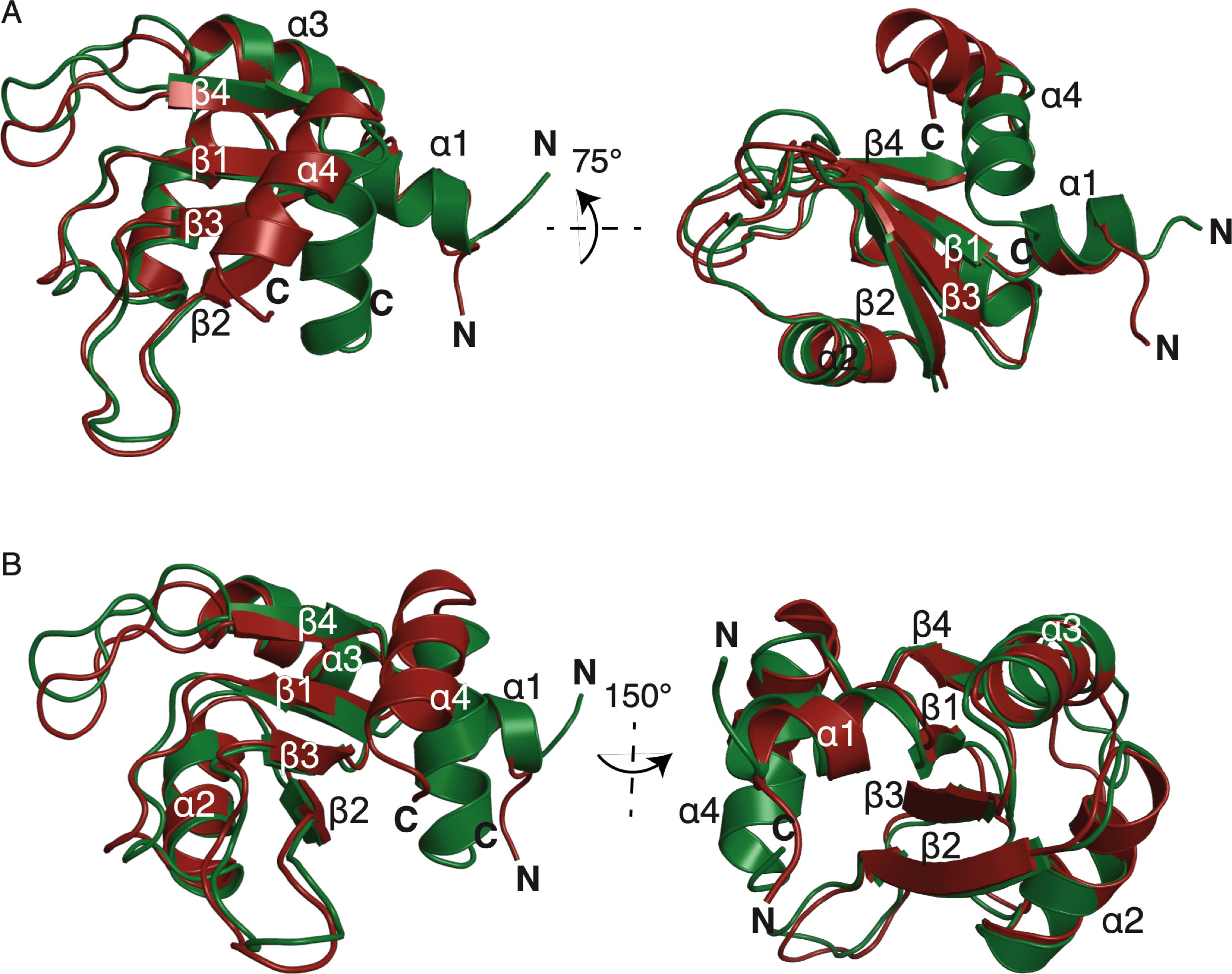

### Supplemental Figure S2

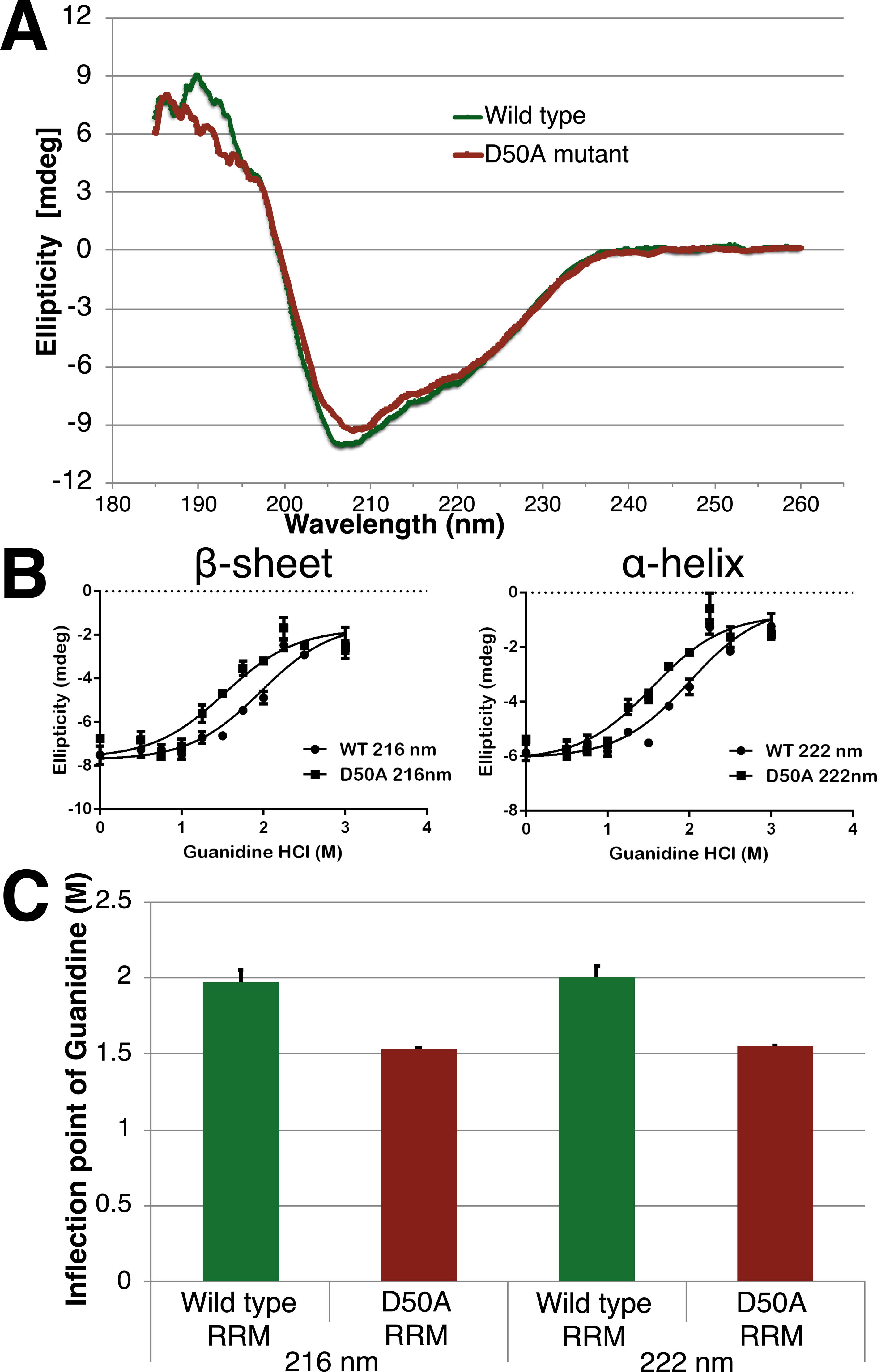

### Supplemental Figure S3

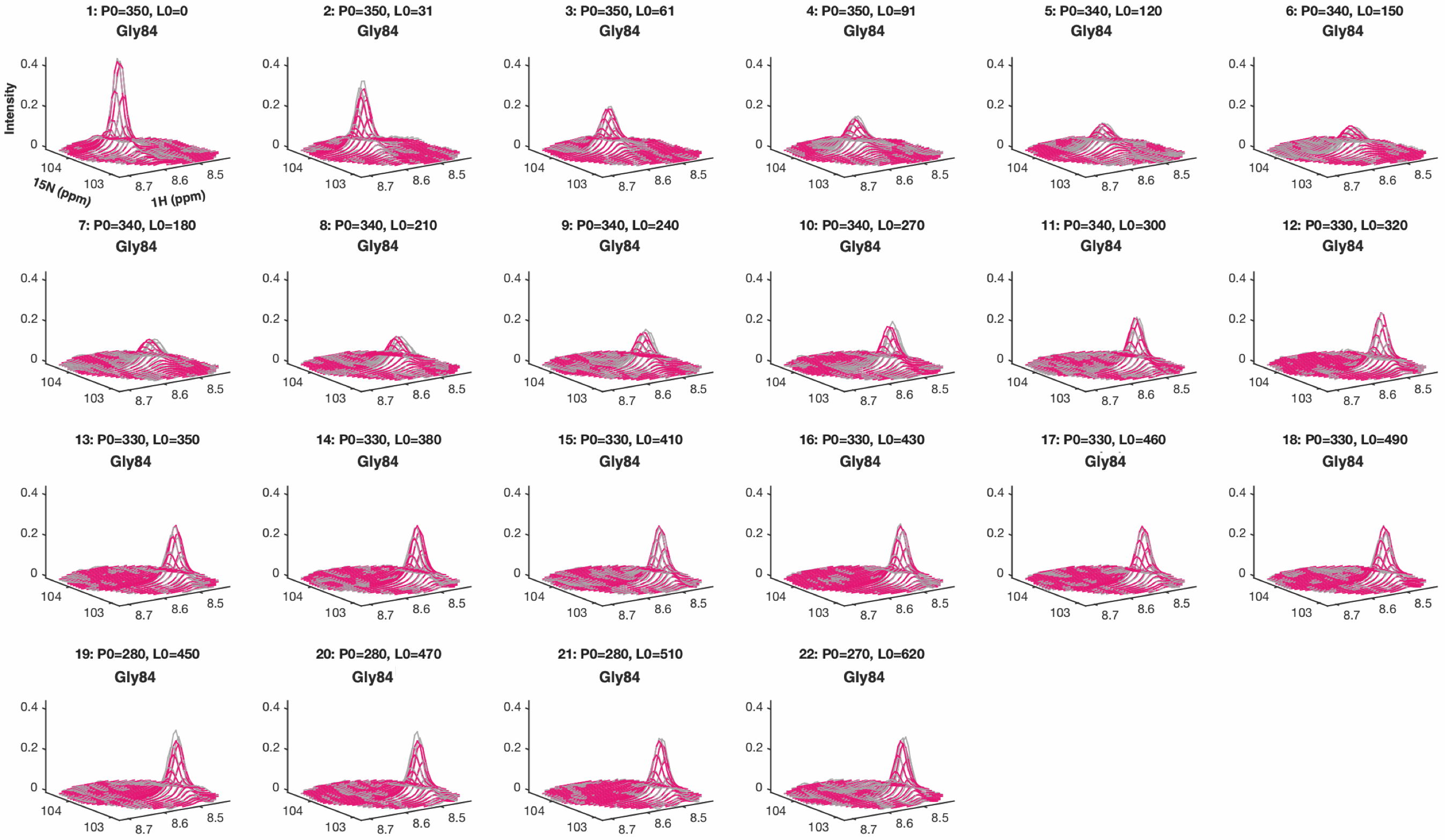

### Supplemental Figure S4

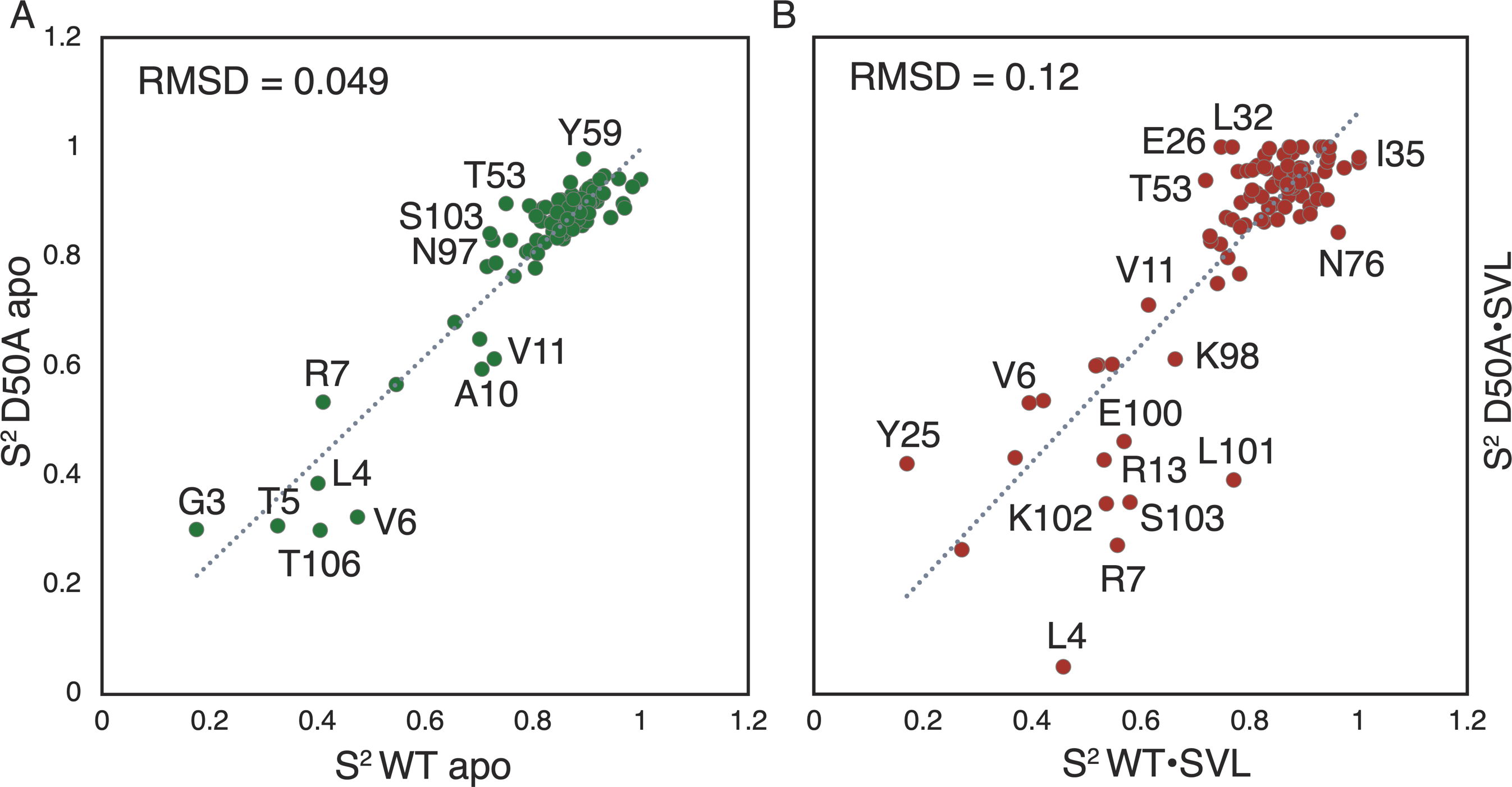

### Supplemental Figure S5

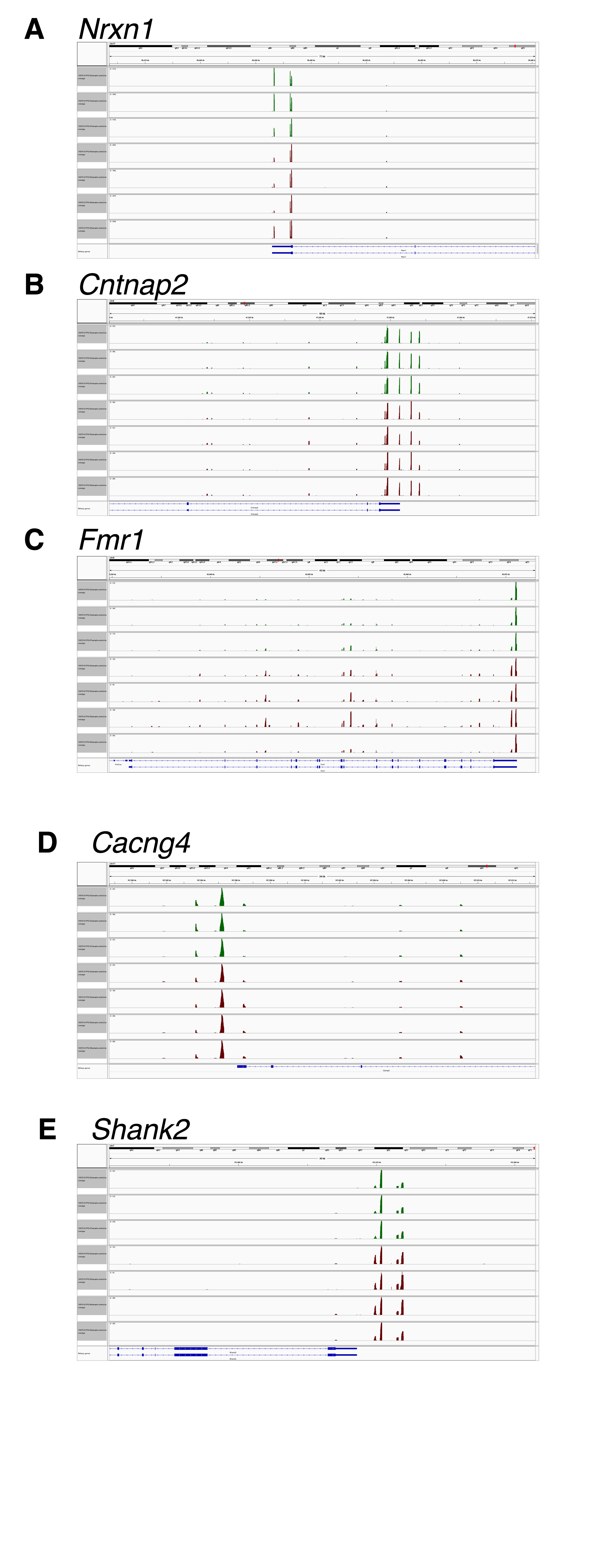
